# A Chromosome-Level Genome Assembly for the Rock Ptarmigan (*Lagopus muta*)

**DOI:** 10.1101/2023.01.31.526508

**Authors:** Theodore E. Squires, Patrik Rödin-Mörch, Giulio Formenti, Alan Tracey, Linelle Abueg, Nadolina Brajuka, Erich Jarvis, Eva C. Halapi, Páll Melsted, Jacob Höglund, Kristinn Pétur Magnússon

## Abstract

The Rock Ptarmigan (*Lagopus muta*) is a cold-adapted, largely sedentary, game bird with a Holarctic distribution. The species represents an important example of an organism likely to be affected by ongoing climatic shifts across a disparate range. We provide here a high-quality reference genome and mitogenome for the Rock Ptarmigan assembled from PacBio HiFi and Hi-C sequencing of a female bird from Iceland. The total size of the genome is 1.03 Gb with a scaffold N50 of 71.23 Mb and a contig N50 of 17.91 Mb. The final scaffolds represent all 40 predicted chromosomes, and the mitochondria with a BUSCO score of 98.6%. Gene annotation resulted in 16,078 protein-coding genes out of a total 19,831 predicted (81.08% excluding pseudogenes). The genome included 21.07% repeat sequences, and the average length of genes, exons, and introns were, 33605, 394, and 4265 bp respectively. The availability of a new reference-quality genome will contribute to understanding the Rock Ptarmigan’s unique evolutionary history, vulnerability to climate change, and demographic trajectories around the globe and serve as a reference genome for the species in the family Tetraonidae (order Galliformes).

## Introduction

The Rock Ptarmigan (*Lagopus muta*, Montin 1776) is a grouse species with a wide distribution across the arctic and subarctic northern hemisphere. It has seasonally variable plumage ranging from almost entirely white in the winter to heavily mottled grey, rust, and brown in the breeding months (see fig. 1). Birds of the genus *Lagopus* are notable for having feathered legs and feet which likely serve to insulate them in cold habitats. The Rock Ptarmigan can be considered as a ring species with variable genetic diversity across its circumpolar range (Sahlman et. al., 2009, Kozma et. al., 2019; see fig. 2). Accordingly, Rock Ptarmigan are expected to be at long-term risk across much of their range due to ongoing climatic changes and limited suitable habitat (Costanzi and Steifetten, 2019; Masanobu et al., 2019).

**Figure 1.**
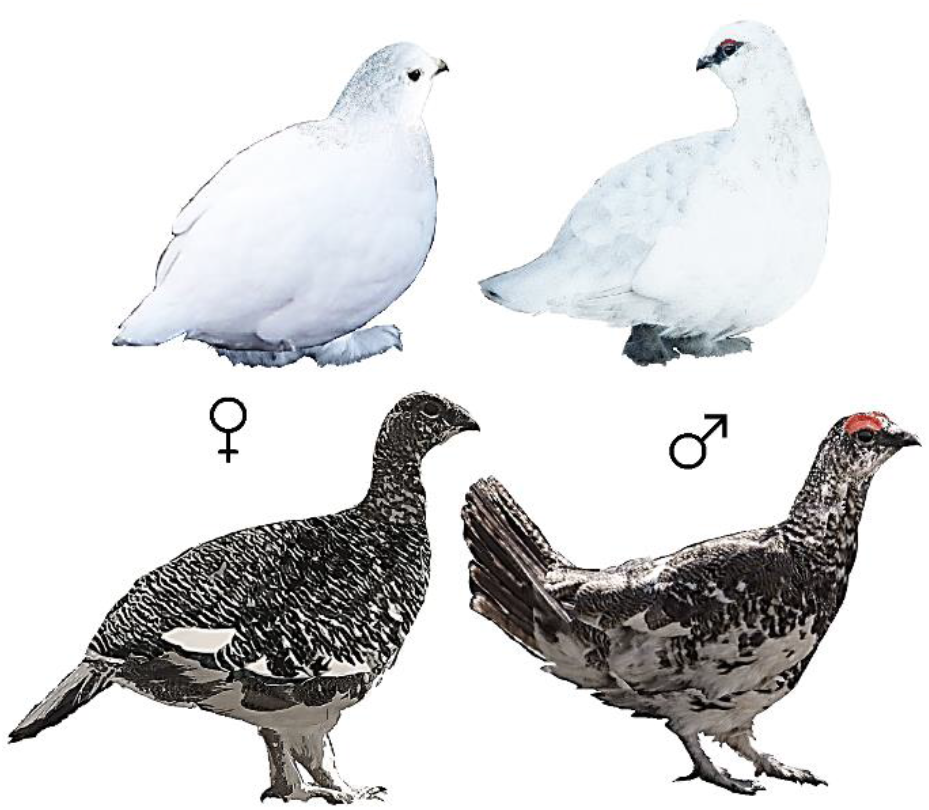
Sexually dimorphic seasonal molt patterns of adult Rock Ptarmigan showing white winter plumage and mottled breeding colors.

**Figure 2.**
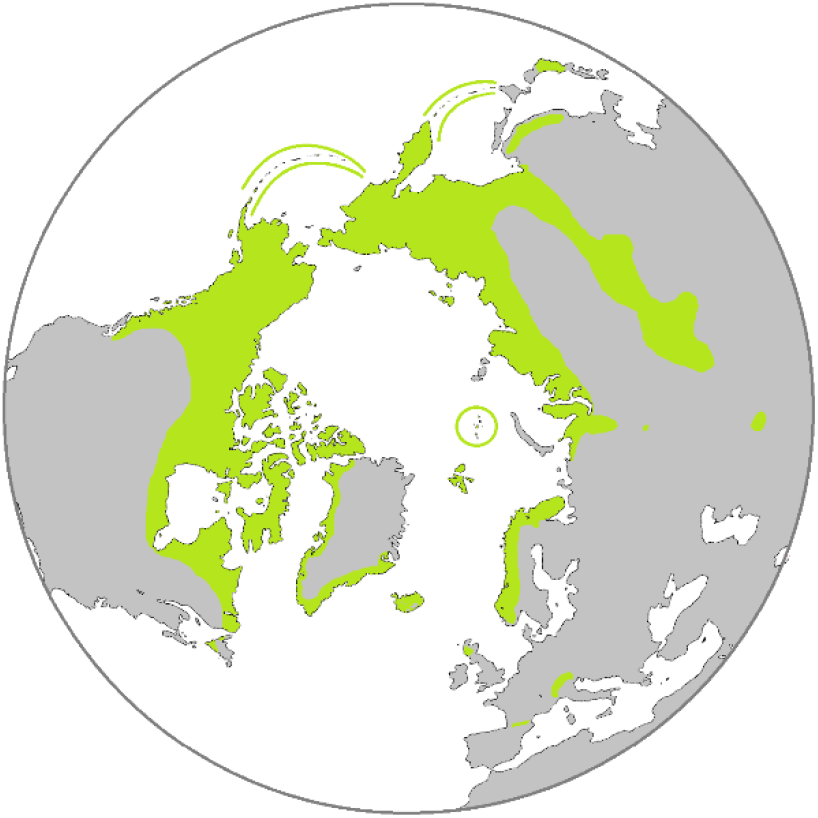
A range map showing the global distribution of Rock Ptarmigan above 30° north.

**Figure 3.**
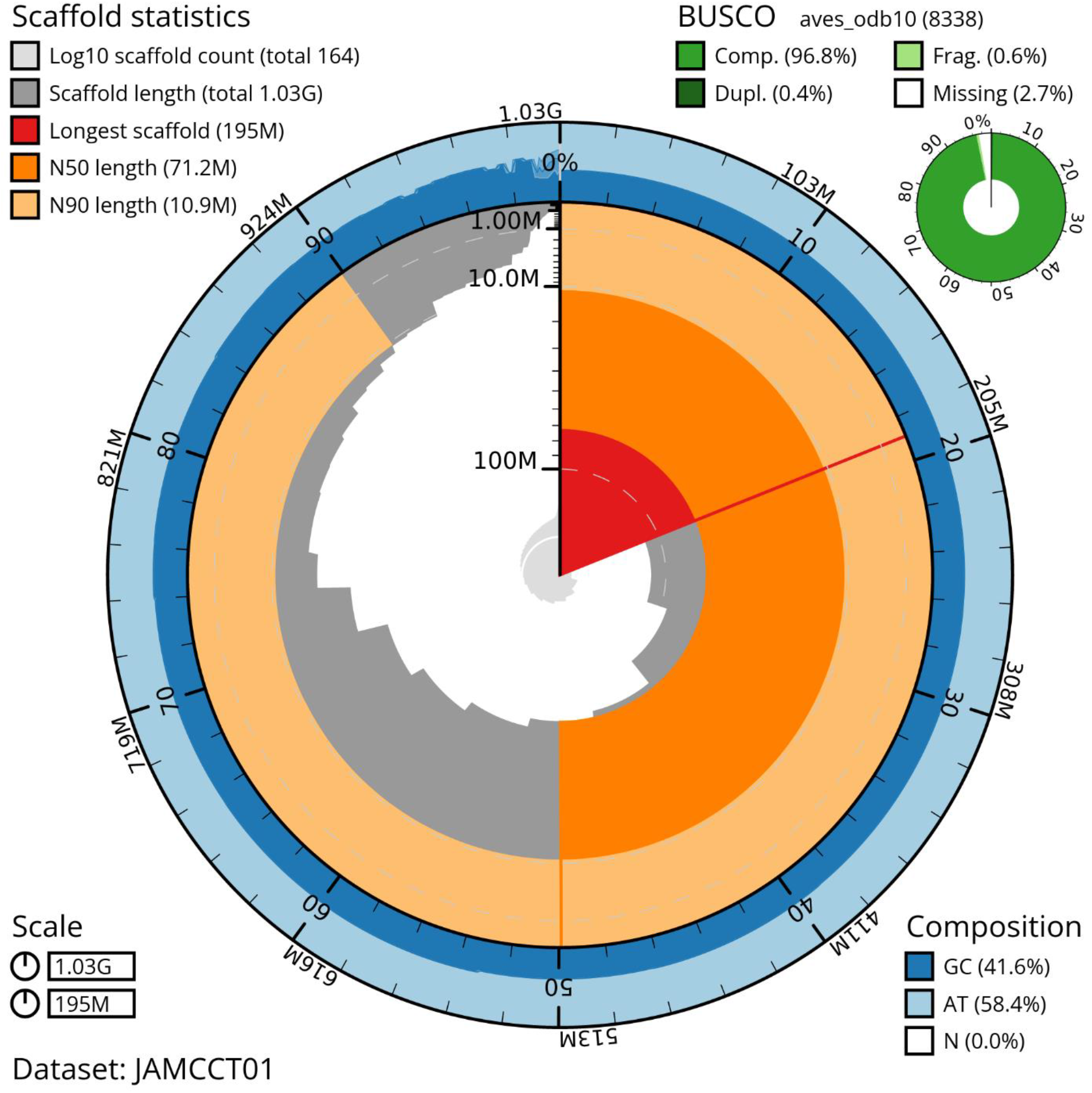
A snail plot indicating the completeness of the bLagMut1 genome assembly. Summary information about scaffold statistics, BUSCO, and the GC vs AT composition of various regions are included.

**Figure 4.**
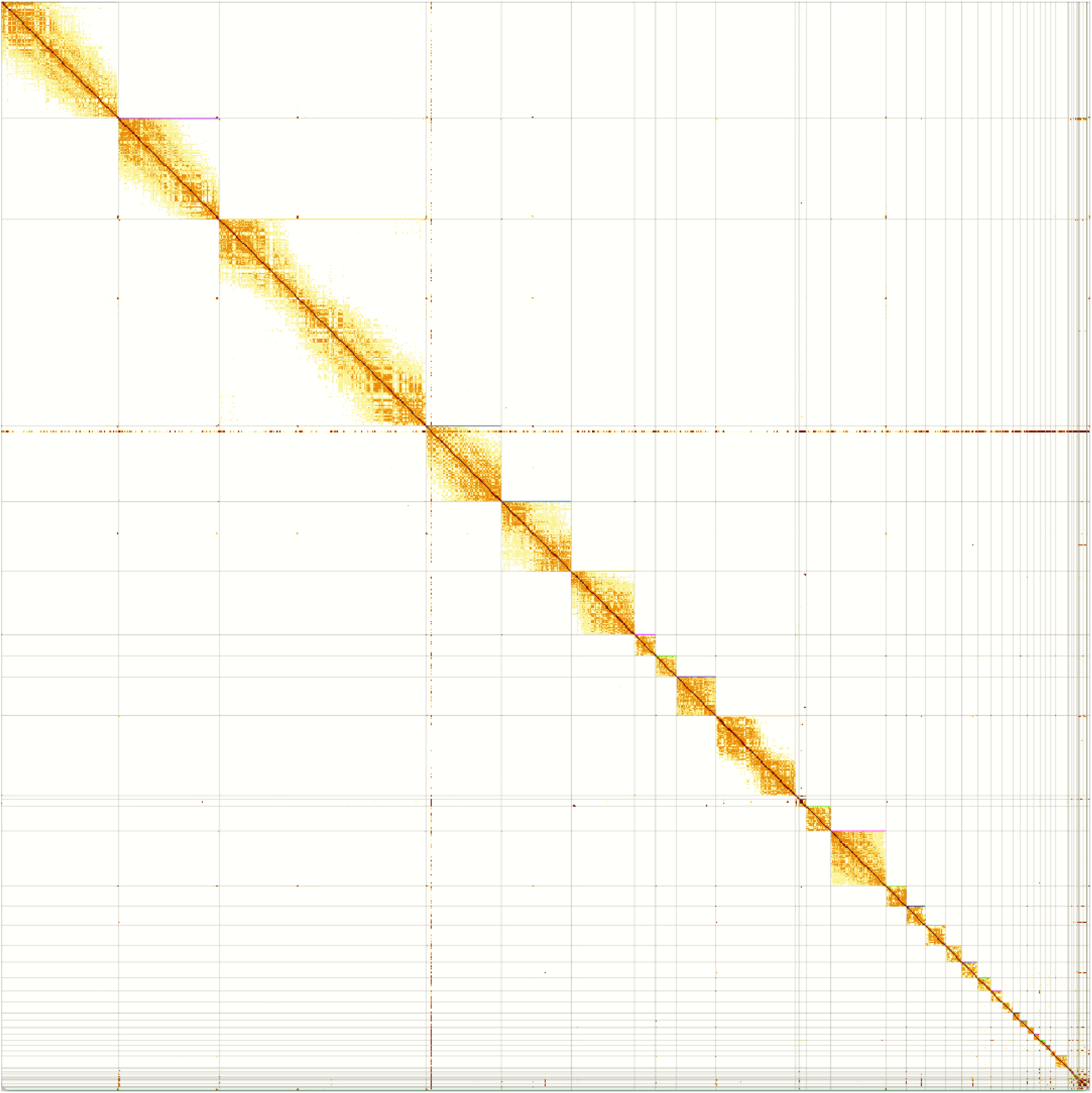
Hi-C contact map for the bLagMut1 genome showing long range contacts generated using PretextView (v. 0.2.5).

With the expected declines in cold specialist species as global temperatures rise (Chamberlain et. al., 2012; Scheffers et al. 2016; Scridel et al., 2019; Höglund et al., 2021), non-migratory birds are particularly valuable to science as they are likely to display many special adaptations necessary for life in the arctic or at high altitude. Some populations of Rock Ptarmigan are considered near-threatened or endangered due to long-term population loss and expected habitat declines (Kozma et al., 2018; Nakamura, 2014; Icelandic Institute of Natural History, 2018). The risks associated with declining genetic quality and environmental changes are not well understood, but might be better assessed with genomic analysis (Bay et al., 2018; Formenti et al., 2022). For populations with robust historical demographics such as the Icelandic Rock Ptarmigan (Gardarsson, 1988; Brynjarsdóttir et al., 2003; Nielsen et al., 1986-2011), a locally sourced reference genome is valuable for assessing demographic history.

The species nearest relatives include other grouse in the subfamily Tetraoninae, although systematics in the order Galliformes remain poorly resolved. The mitochondrial genome of Rock Ptarmigan was previously made available along with the mitochondrial DNA of a sister species Willow Grouse (*Lagopus lagopus*; Sveinsdóttir and Magnússon, 2017). The Willow Grouse and Rock Ptarmigan are believed to have diverged as recently as 2-5 million years ago (Persons et al., 2016) and are often studied together (Kozma et al., 2018; Lucchini et al., 2001). The white-tailed ptarmigan (*Lagopus leucura*) is the most closely related species with whole genome data available, having a common ancestor with other *Lagopus* taxa no older than 3 million years ago, although the genome assembly is not currently annotated (Kozma et al., 2019; Clark et. al., 2016; GenBank: GCA_019238085.1).

Here, we describe the first reference-quality genome assembly and annotations for Rock Ptarmigan. A combination of long-read and conformation capture sequencing technologies were used to assemble a 1.03 Gb haploid reference genome.

## Materials and methods

### Sample collection and PCR preparation

As basis for the reference genome assembly and annotation, fresh blood from a single female bird collected (shot) in Húsavík, northern Iceland, in 2018 was used (NCBI BioSample SAMN25144835) while additional, heart, muscle, brain, kidney, liver, ovaries, testes, and spleen from a second bird was collected for RNA-seq to aid in gene prediction (NCBI BioSample SAMN26436951, SAMN29421920, SAMN29421921, SAMN29421922, SAMN29421923, SAMN29421924, SAMN29421925, and SAMN29421926 respectively). DNA extraction was performed in the laboratories of SciLifeLab (Uppsala, Sweden). RNA was isolated, at University of Akureyri, using Beckman Coulter RNAClean XP (FisherScientific, USA). Materials from the birds used for the genome assembly are stored at the Icelandic Institute of Natural History in Garðabær, Iceland (Accession nr. RM13211).

### Sequencing

Whole genome sequencing was carried out using two PacBio SMRT smart cells run on a Sequel II system at SciLifeLab in Uppsala, while Dovetail Genomics Hi-C Kits were processed on an Illumina NovaSeq 6000 at SciLifeLab in Stockholm. RNAseq was carried out on Illumina HiSeq2500 system Paired-end 2×125 cycles at deCODE genetics, Reykjavík.

### Genome assembly

The genome was assembled following the Vertebrate Genome Project (VGP; Rhie, 2021) assembly pipeline. First, a kmer database was generated using Meryl (v. 1.3) from the PacBio HiFi reads for reference-free genome evaluation and downstream assembly QC. The kmer size was set to 21 after running the best_k.sh script for the expected genome size (~1Gb) in Merqury (v. 1.3; Rhie et al., 2020). PacBio HiFi reads were assembled using hifiasm (v. 0.15.1-r334; Cheng et al., 2021), followed by a round of purge_dups (v. 1.2.5; Guan et al., 2020) incorporating minimap2 (v. 2.17-r941). Each of the previous steps was followed by assembly evaluation. This included contig/scaffold statistics computed using the Python library assembly_stats (v. 0.1.4), and BUSCO (v. 5.3.1), while completeness and quality value statistics of the assembly along with kmer spectrum plots were produced using Merqury (v. 1.3; Rhie et al., 2020). The assembly was scaffolded using the Hi-C reads. Briefly, reads were first aligned to the assembly using the VGP modified version of the Arima mapping pipeline that uses bwa mem (v. 0.7.17-r1188) and samtools (v. 1.19) for alignment and Picard (v. 2.10.3) for 5’ end filtering and duplication removal. Scaffolding was performed using Salsa2 (v. 2.3) and evaluated using BUSCO and scaffold statistics.

The Hi-C reads were then mapped back to the scaffolded assembly using the same pipeline as in the previous step and the resulting bam file was converted to pretext format using PretextMap (v. 0.1.7).

The finalized assembly was screened for contamination and then manually curated (Howe et al., 2021). Curation was performed using gEVAL (Chow et al., 2016) and Hi-C contact maps visualized in HiGlass (Kerpedjiev et al., 2018) and PretextView (v. 0.2.5), resulting in 97 missed or mis-join corrections to the scaffolds producing a resolved chromosome level genome with 38 autosomes and, the Z and W sex chromosomes. Construction of microchromosomes was investigated using the Mummer alignment tool (v 4.0.0: Marçais et al., 2018) although poor syntany was noted for comparison with Gallus gallus and some likely remain unresolved.

The mitochondrial genome was assembled separately from both raw reads and contigs using MitoHifi (v. 2.2; Uliano-Silva et. al., 2021) with automatic alignment to the Japanese Rock Ptarmigan (*L. muta japonica;* Yonezawa and Nishibori, 2020) via built-in features from the MitoFinder dependency (v. 1.4.1; Allio et. al., 2020).

The completed genome assembly is publicly available in NCBI under accession number GCA_023343835.1. The mitochondrial assembly is publicly available in NCBI under BankIt submission 2677809.

### Genome annotation

The Rock Ptarmigan reference genome was annotated using the standard NCBI Eukaryotic Genome Annotation Pipeline version 10.0. A detailed summary of the pipeline is available online at: https://www.ncbi.nlm.nih.gov/genome/annotation_euk/process/. In contrast to previous iterations, this version of the pipeline used RFAM (v. 14.6; Kalvari et al., 2021) for discovery of small non-coding RNA’s and STAR (Dobin et al. 2013) for alignment of RNA-seq reads from our supplementary tissues. The pipeline has stable use of several tools including BUSCO (v. 4.1.4; Manni et al., 2021) and Splign (Kapustin et al., 2008) among others.

For calculation of genomic masking, the Rock Ptarmigan genome was masked with WindowMasker (Morgulis et al., 2006). Annotation of the mitochondrial genome was achieved via manual comparison with the extant published Icelandic Rock Ptarmigan mitogenome in addition to automatic annotation using MITOS WebServer (Bernt et. al., 2013).

## Results

### Sequencing and Assembly Results

The final assembly sequence is 1,026,771,810 base pairs long, with 71,937 gap bases (0.007%) across 210 spanned gaps. The genome assembly includes 375 contigs arranged on 165 scaffolds. The scaffold N50 is 71,229,700 bp with an L50 of 5. The Contig N50 is 17,905,263 bp with an L50 of 19.

Average coverage across the genome is 57.75x. In total 38 autosomes were identified, with 18 unlocalized sequences among them. Additional W and Z allosomes were described with only a single unlocalized sequence found on the W. Assembly summary statistics appear significantly better than the current *Gallus gallus* reference genome (GRCg6a), and are modest in comparison to the most recently annotated *Gallus gallus* individual (bGalGal1.mat.broiler.GRCg7b; See Table 1 below). Kmer spectra plots overall showed the expected copy-kmer distributions (See Figure 5 Below).

**Table 1.**
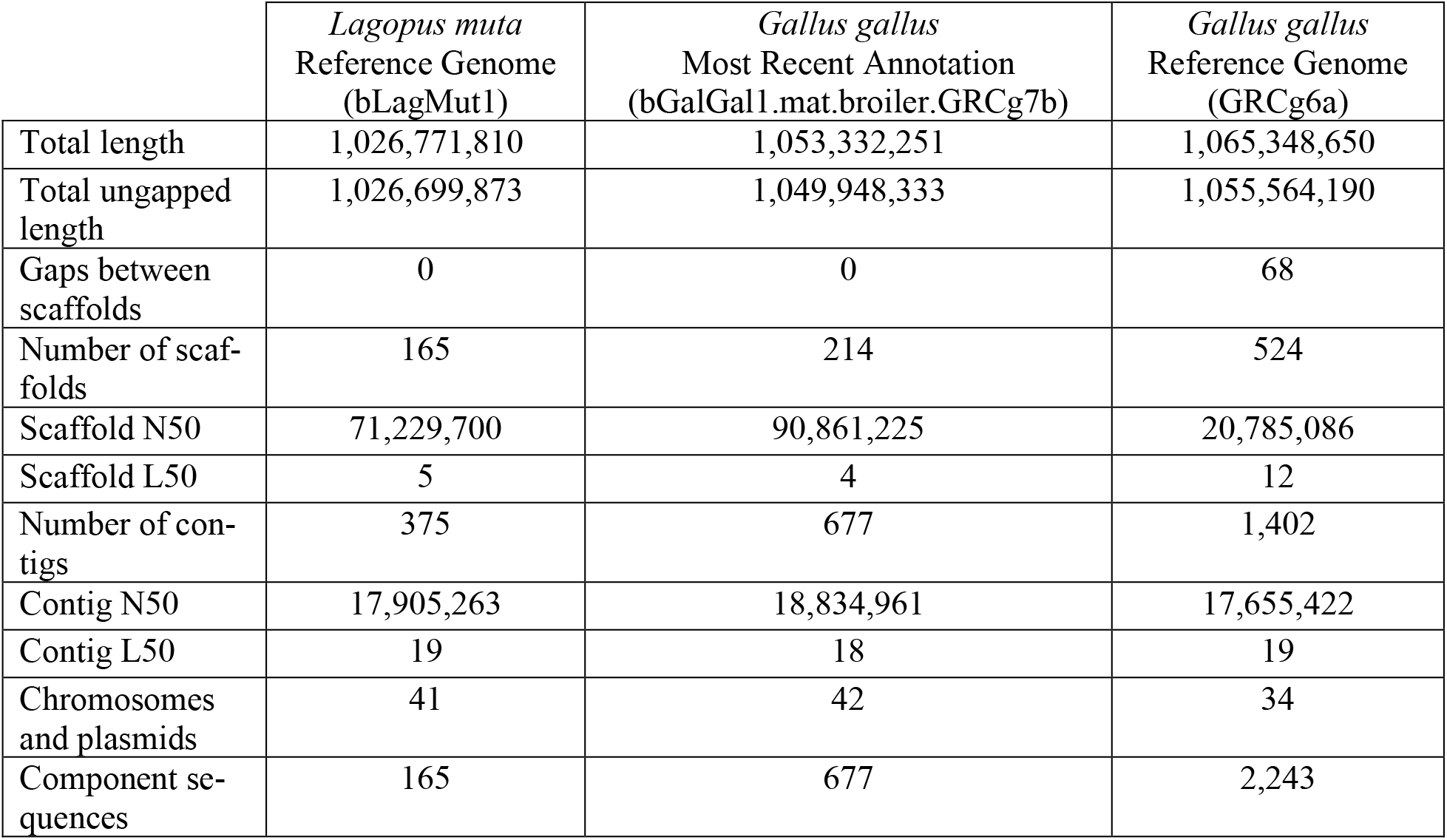
A series of “global” statistics published in the public release of the Rock Ptarmigan reference genome on NCBI indicating the completeness of the new reference genome in comparison to the gold standard Chicken reference genome and the most recently annotated Chicken reference genome.

**Figure 5.**
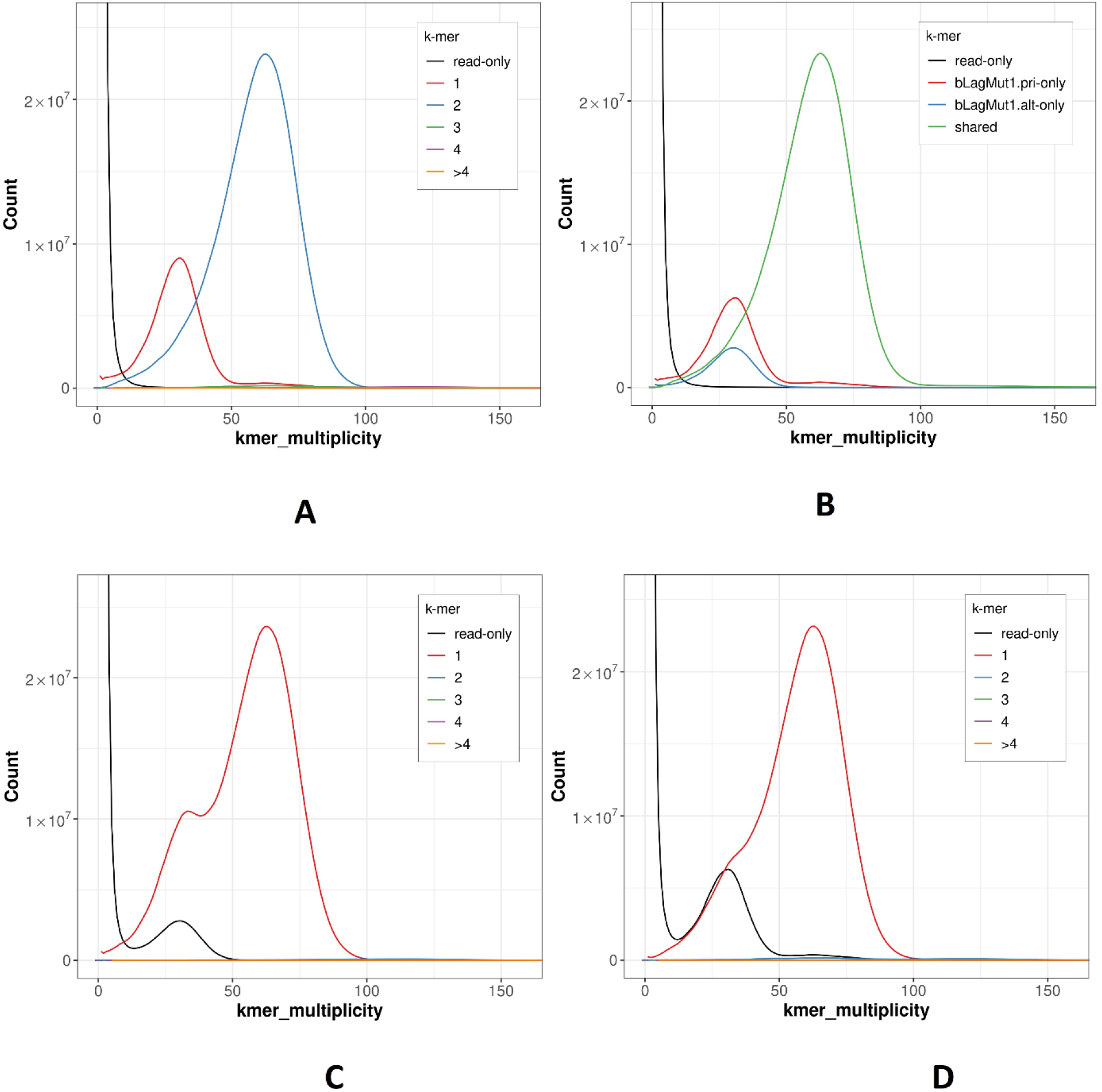
Outputs from Merqury showing kmer distribution according to: (A.) Spectra-cn plot of the bLagMut1 complete assembly, (B.) spectra-asm plot of the bLagMut1 complete assembly, (C.) spectra-cn plot of the bLagMut1 primary assembly, and (D.) spectra-cn plot of the bLagMut1 alternate assembly.

### Genome Annotation

In total 20,110 genes and pseudogenes were identified by combining gene prediction and similarity approaches, with approximately 80% identified as protein coding. The annotated genes showed a 98.6% completeness score against 98.9% for the whole genome when set against the BUSCO avian dataset (aves_odb10 lineage) and indicating 0.9% of genes missing from the annotated assembly. The annotation and associate summary statistics are available in NCBI’s RefSeq genome record for the reference (Pruitt et al., 2013). The contents of the report are summarized in Table 2.

**Table 2.** Comparative table showing the relative accuracy and completeness of the *Lagopus muta* reference annotation (NCBI *Lagopus muta* Annotation Release 100) against the most recently complete annotation of the chicken genome (NCBI *Gallus gallus* Annotation Release 106).

|  | *Lagopus muta*<br>Reference Genome Annotation<br>(bLagMut1) | *Gallus gallus*<br>Most Recent Annotation<br>(bGalGal1.mat.broiler.GRCg7b) |
| --- | --- | --- |
| Genes and Pseudogenes | 20,110 | 25,635 |
| Protein Coding Genes | 16,078 | 18,023 |
| Non-Coding Genes | 3,738 | 7,330 |
| mRNA | 43,785 | 68,670 |
| lncRNA | 5,431 | 10,062 |
| tRNA | 306 | 303 |
| CDSs | 43,793 | 68,683 |
| Introns (mean length) | 206,142 (4,265) | 241,290 (4,145) |
| Exons (mean length) | 229,018 (394) | 262,919 (490) |
| Mean Gene Size | 33kb | 28kb |
| Maximum Gene Size | 1.6 Mb | 1.3 Mb |
| BUSCO Score | 98.6% | 98.7% |

### Mitochondrial Genome

The mitochondrial DNA was described with all 13 expected protein coding regions and analyzed for accuracy through comparative analysis. With our addition, there are now four extant mitochondrial genomes published for the Rock Ptarmigan; Two from Iceland, one from Japan, and one from Siberia (Sveinsdóttir and Magnússon, 2017; Yonezawa and Nishibori, 2020; Wang et. al., 2017). Using the ClustalW package embedded in BioEdit (Thompson et. al., 1994; Hall, 1999), we found a total of 24 bases divergent from the previously published Icelandic Rock Ptarmigan mitogenome in a manual review. Of these divergences 14 appeared in coding regions and 8 appeared unique to the previously published individual and our calls at those locations were conserved in the other Rock Ptarmigan populations. None of the polymorphisms observed between the populations appeared to be uniquely conserved in the Icelandic Population. Analysis of pairwise distances using phylogenetic tree software in Mega11 (Tamura et. al., 2021) showed clear grouping of the Rock Ptarmigan separated from the Willow Ptarmigan, as previously reported (Sveinsdóttir and Magnússon, 2017).

## Discussion/Conclusion

Our avian reference genome includes a highly complete set of information with 99.994% of the 1.03 Gb described matching to 40 haploid chromosomes and the mitochondria. Other recent works have aimed to unlock the potential provided by Rock Ptarmigan genetics (Kozma et al., 2018; Kozma et al., 2019; Sigmarsdóttir, 2022). As observed for other recently published genomes (Formenti et al., 2022), the new Rock Ptarmigan genome is of comparatively excellent quality (see also Table 1).

Though Rock Ptarmigan has been globally identified as Least Concern by the IUCN in recent years, there have been regional fluctuations in its status and some nations identify the species as threatened due to long term declines (IUCN, 2022; European Commission, 2022; Icelandic Institute of Natural History, 2018). There is evidence that sub-populations of other grouse species may poses important local adaptations necessary for persistence (Oh et al., 2019), making it probable that the Rock Ptarmigan has unique evolutionary adaptations across its range. Further, it is well established that Arctic species such as Rock Ptarmigan may be disproportionately affected by climate change with an expected poleward contraction of species’ ranges (Birdlife International, 2015; Kozma et al., 2018). For more disparate populations such as those in the Japanese mountains of Honshu, the European Alps, and the Pyrenees, rising tree lines may entirely squeeze the Rock Ptarmigan out of its montane niches as has been suggested broadly for alpine habitats (Dirnböck et al., 2011; MRI EDW Working Group, 2015), and some closely related species (Jackson et al., 2016). In the context of conservation, having a reference genome available will contribute to our understanding of the species’ genetic risks and possible movements in the face of a warming planet (Bay et al., 2018; Kozma et al., 2018).

Many wildlife species are difficult to study at the genomic level due to limited specimen availability and constraints on procurement (Hope et al., 2018; Kemp, 2015). Because the Rock Ptarmigan is a widespread game bird, it is particularly useful for both genomic studies and general investigations into wildlife ecology. Hunters have the potential to contribute robust data regarding the species trends and may continue to contribute both historical and new specimen materials for research (Cretois et al., 2020). Given the species’ close cultural connection to some regions and history as a food source (McGovern et al., 2006), the Rock Ptarmigan may benefit from additional conservation efforts from an involved public or concerned hunters and may be a good candidate for flagship status (McGowan et al., 2020).

Future studies into Rock Ptarmigan genomics will benefit from decades of studies into these birds in captivity (Stokkan et al., 1988). Recently, Rock Ptarmigan hatched and raised in captivity have been used for gene expression studies to understand circadian rhythms and investigate the cecal microbiome representing valuable opportunities going forward (Appenroth et al., 2021; Appenroth et al., 2020; Salgado-Flores et al., 2019).

Among avian diversity, the birds in the family Galliformes represent less than 3% of all species but have an outsized impact on global economics with Chickens, Turkeys, Pheasants, Quails, and Grouse all being regularly consumed. Among the available avian genomes (Bravo et al., 2021) those in order Galliformes are represented with 26 species assemblies currently available on NCBI (approximately 5% of all extant; Sayers et al., 2022). Among these, 68 assemblies have been completed and the chicken has been assembled 30 times (for context see Burt, 2005; Li et al., 2022). This highlights a commercial implication for Rock Ptarmigans as they have many special adaptations that could be of importance to domestic poultry.

Given the usefulness of wild relatives for research into domesticated species (Li et al., 2020; Jackson et al., 2016) the Rock Ptarmigan may prove to be a useful model for understanding other Galliformes. This relationship will surely have limitations in the genomic realm as more distantly related species are less informative at finer scales than those that are closely related (Scutari et al., 2016). However, if the Rock Ptarmigan’s genes tailored to arctic landscapes can be used to better understand genetic architecture for cold weather survival, improved forage capabilities, or other ancestral traits, then important pathways may be identified for commercially exploited birds or other species of conservation interest.

Taking all of this into consideration, the availability of a Rock Ptarmigan reference genome makes the species exceptionally well positioned for investigation across a broad new range of scientific inquiry. With links to arctic/alpine biomes, conservation, hunting culture, and industry, the Rock Ptarmigan reference genome provides a unique opportunity to investigate a species at the intersection of many issues of global significance.

## Significance statement

The Rock Ptarmigan is a widespread bird species of economic and nutritional importance to large portions of the northern hemisphere. Only a tiny fraction of the Rock Ptarmigan’s genome was previously reported and studied. The effort undertaken to sequence and annotate the whole genome provides an ability to understand the species at a molecular level. This vertebrate genome allows for new critical assessment of the Rock Ptarmigan and related species at the individual, population, and environmental scales.

## Acknowledgments

We wish to express our thanks to Ignas Bunikis at SciLifeLab, Uppsala, for assistance with the assembly and scaffolding work. We also wish to extend our appreciation to Ólafur Þ. Magnússon at deCODE genetics, Reykjavik, for arranging RNAseq.

## Data Availability Statement

The final annotation has been publicly released and uploaded according to the high standards of the Earth BioGenome Project (Lewin et al., 2018). The genome assembly, including the raw shotgun sequencing data, and the mitochondrial genome has been uploaded to NCBI and is available at https://www.ncbi.nlm.nih.gov/assembly/GCA_023343835.1; BioProject: PRJNA836583; BioSample: SAMN25144835

